# Repurposing cytarabine for treating primary effusion lymphoma by targeting KSHV latent and lytic replications

**DOI:** 10.1101/299339

**Authors:** Marion Gruffaz, Shenghua Zhou, Karthik Vasan, Teresa Rushing, Qing Liu Michael, Chu Lu, Jae U. Jung, Shou-Jiang Gao

## Abstract

Oncogenic Kaposi’s sarcoma-associated herpesvirus (KSHV) is etiologically linked to primary effusion lymphoma (PEL), an aggressive and non-treatable malignancy commonly found in AIDS patients. In this study, we performed a high throughput screening of 3,731 characterized compounds, and identified cytarabine approved by FDA for treating numerous types of cancer as a potent inhibitor of KSHV-induced PEL. We showed the high efficacy of cytarabine in the growth inhibition of various PEL cells by inducing cell cycle arrest and apoptosis. Cytarabine inhibited host DNA and RNA syntheses and therefore induced cellular cytotoxicity. Furthermore, cytarabine inhibited viral DNA and RNA syntheses and induced the the rapid degradation of KSHV major latent protein LANA, leading to the suppression of KSHV latent replication. Importantly, cytarabine effectively inhibited active KSHV replication and virion production in PEL cells. Finally, cytarabine treatments not only effectively inhibited the initiation and progression of PEL tumors, but also induced regression of grown PEL tumors in a xenograft mouse model. Together, our study has identified cytarabine as novel therapeutic agent for treating PEL as well as eliminating KSHV persistent infection.

**Importance:** Primary effusion lymphoma is an aggressive malignancy caused by Kaposi’s sarcoma-associated herpesvirus. The outcome of primary effusion lymphoma is dismal without specific treatment. Through a high throughput screening of characterized compounds, we identified a FDA-approved compound cytarabine as a potent inhibitor of primary effusion lymphoma. We showed that cytarabine induced regression of PEL tumors in a xenograft mouse model. Cytarabine inhibited host and viral DNA and RNA syntheses, resulting in the induction of cytotoxicity. Of interest, cytarabine induced the degradation of KSHV major latent protein LANA, hence suppressing KSHV latent replication, which is required for PEL survival. Furthermore, cytarabine inhibited KSHV lytic replication program, preventing virion production. Our findings identified cytarabine as novel therapeutic agent for treating PEL as well as for eliminating KSHV persistent infection. Since cytarabine is already approved by the FDA, it might be an ideal candidate for repurposing for PEL therapy and for further evaluation in advanced clinical trials.

## Introduction

Kaposi’s sarcoma-associated herpesvirus (KSHV) is an oncogenic gammaherpesvirus displaying a biphasic lifecycle of latent and lytic replication phases (1). Following primary infection, KSHV establishes a lifelong latent phase punctuated by reactivation into lytic replication. KSHV is associated with several malignancies including Kaposi’s sarcoma (KS), primary effusion lymphoma (PEL), a subset of multicentric Castleman’s disease (MCD), and KSHV-associated inflammatory cytokine syndrome (KICS) (2).

PEL is a B-cell neoplasm involving body cavities of pleural, pericardial and peritoneal spaces usually without extracavitary tumor masses (3). All PEL cells harbor multiple copies of KSHV genome, which are required for their survival. Up to 70% of PEL cases are also associated with Epstein-Barr virus (EBV) infection (3). Most PEL cells are latently infected by KSHV but a small number of them undergoes spontaneous lytic replication. Several KSHV latent genes including LANA (ORF73), vCyclin (ORF72), vFLIP (ORF71) and a cluster of microRNAs (miRNAs) drive the proliferation and survival of PEL cells (3, 4). Numerous viral lytic genes such as vIL6 (ORF-K2) also play a role in PEL growth and survival (3).

PEL usually occurs in HIV-infected patients, of which half have KS or a history of KS (3). PEL accounts for about 4% of non-Hodgkin’s lymphomas (NHLs) in HIV patients (5, 6). Rare cases of PEL have been described in HIV-negative immunocompromized patients after solid transplantation, or elderly men living in areas with a high KSHV prevalence such as Mediterranean and Eastern European regions (7, 8).

The prognosis of PEL patients is usually poor with a median survival time of 6.2 months (9). There is currently no efficient and specific treatment for PEL (2, 3). Because of its rarity, there has been so far no large prospective clinical trial to investigate the proper therapy for PEL. Hence, finding a treatment for PEL remains a challenge. In this context, repurposing old drugs is an attractive strategy for identifying potential treatment options for PEL.

Here, we performed a high throughput screening (HTS) of 3,731 characterized compounds to identify inhibitors targeting KSHV-induced oncogenic addiction and malignancies. We used a model of KSHV-induced cellular transformation of rat primary mesenchymal stem cells. This model offers both parallel uninfected (MM) and transformed cells (KMM) for comparative screening (10). We identified cytarabine as a promising candidate for targeting KSHV-induced oncogenic addiction. Cytarabine is currently approved by FDA for treating acute myeloid leukemia (AML), acute lymphocytic leukemia (ALL) and NHLs (11). We demonstrated that cytarabine was effective in inducing cell cycle arrest and apoptosis in PEL cells. Furthermore, cytarabine effectively inhibited the initiation and progression of PEL, and regressed grown PEL tumors in a xenograft mouse model. Importantly, cytarabine not only did not trigger, rather it inhibited KSHV lytic replication program, preventing virion production. Mechanistically, cytarabine inhibited both cellular and viral DNA and RNA syntheses, and triggered the degradation KSHV major latency-associated nuclear antigen (LANA), hence inducing cell stress and inhibiting KSHV persistent infection.

## Results

### Identification of inhibitors of KSHV-transformed cells

To identify inhibitors of KSHV-transformed cells, we conducted a HTS with libraries of small molecules using MM and KMM cells (10). The libraries consist of 3,731 individual compounds, including the 2,320 compounds of Spectrum collection from MicroSource Discovery System Inc. covering a wide range of biological activities and structural diversities that are suitable for HTS programs; the NIH Clinical collection consisting 781 compounds previously tested in clinical trials; the EMD Millipore Kinase collection consisting of 327 well-characterized, pharmacologically active, potent protein kinase and/or phosphatase inhibitors; and the EMD Millipore StemSelect small molecule regulators collection consisting of 303 pharmacologically active compounds including extracellular domain-targeting reagents as well as cell-permeable reagents that regulate intracellular targets. We treated MM and KMM cells with 5 µM of each compound for 48 h and counted the surviving cells following staining with DAPI. We selected 50 compounds representing 1.3% of the libraries that induced cytotoxicity in >50% of KMM cells and <10% of MM cells (Fig.1A and B). Interestingly, most of the selected compounds (54%) are anti-inflammatory, followed by antibacterial (10%), antihypertensive (10%), antioxidant (4%), antiviral (2%), antiarthritic (2%), antiangiogenesis (2%) and cathecholaminegic (2%) (Fig. 1C). Among them, cytarabine, a cytidine analogue with a modified sugar moiety, *i.e*.arabinose instead of ribose, is currently used for treating leukemia and lymphomas (11), and hence is an interesting candidate that could be repurposed for KSHV-induced malignancies.

**FIG 1.**
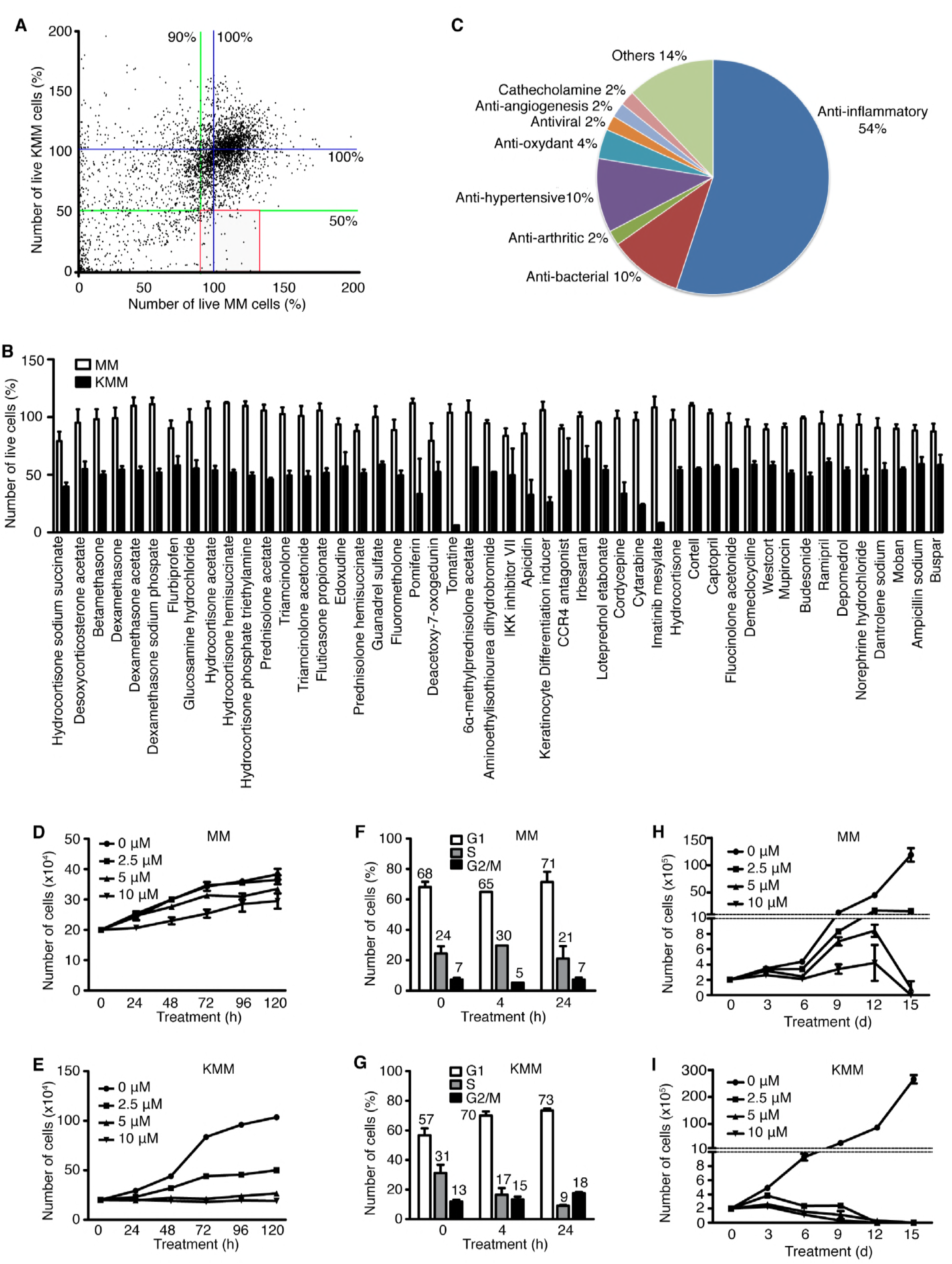
Identification of small molecules that induce cytotoxicity in KSHV-transformed (KMM) but not uninfected (MM) cells. (A) MM and KMM cells were treated with libraries of 3,731 compounds from the MicroSource Discovery System Spectrum collection, the NIH Clinical collection, the EMD Millipore kinases collection and the EMD Millipore StemSelect small molecule regulators collection. Cells treated with 5 μM of each compound for 48 h were washed with PBS, fixed, stained with DAPI and counted for live cells. Results normalized using DMSO as a negative control (standardized at 100% of live cells) were expressed as percentages of live cells. The blue lines represent 100% live cells. The green lines are cutoff thresholds set at 50% of live cells for KMM cells and 90% of live cell for MM cells. A total of 73 compounds (1.9% of all compounds) shown in the red square were selected from the initial screening. (B) Secondary screening to validate the 73 selected compounds resulting in the selection of 50 compounds (1.3% of all compounds). Error bars represent the pool of the results from the first screening and the secondary screening. (C) Analysis of the biological functions of the 50 validated compounds. Results are expressed in percentages. (D-E) Analysis of the effect of cytarabine on cell proliferation of MM (D) and KMM (E) cells. (F-G) Analysis of the effect of 5 μM cytarabine on cell cycle progression of MM (F) and KMM (G) cells. (H-I) Analysis of the long-term effect of different doses of cytarabine on MM (H) and KMM (I) cells over a period of 15 days. Cytarabine was replenished every 3 days.

To confirm the inhibition efficacy of cytarabine on KSHV-transformed cells, we treated MM and KMM cells with 0, 2.5, 5 and 10 µM of cytarabine over a period of 120 h. Cytarabine preferentially inhibited the proliferation of KMM cells in a dose-dependent manner (Fig. 1D and E). Consistently, cytarabine induced cell cycle arrest in KMM but not MM cells (Fig. 1F and G). Furthermore, extended treatment with cytarabine for up to 15 days completely eliminated live KMM cells by day 9 at 10 µM and by day 12 at both 2.5 and 5 µM (Fig. 1I). In contrast, while cytarabine at 5 and 10 µM had slight inhibitory effect on MM cells, most cells survived up to day 12; however, most of the cells detached by day 15 albeit they remained alive (Fig. 1H).

### Cytarabine inhibits the proliferation of diverse PEL lines

We tested the inhibitory efficacy of cytarabine on different PEL lines including BCBL1, BC3, JSC1 and BCP1 cells that are singly infected by KSHV, and BC1 cells that are dually infected by KSHV and EBV. As there is no appropriate control for PEL cells, we included BJAB, a KSHV-and EBV-negative Burkitt’s lymphoma cell line as a reference. At 0.5, 2 and 5 µM, cytarabine effectively inhibited the proliferation of all PEL lines tested, which manifested a greater sensitivity than BJAB cells (Fig. 2A-F). As a result, PEL lines had IC50 ranging from 0.44 µM to 1.29 µM while BJAB cells had an IC50 2.34 µM (Fig. 2G). Extended treatment of BCBL1 and BCP1 cells with 1 µM cytarabine for up to 15 days completely killed all cells of both lines by day 9, indicating the lack of any emerging resistance (Fig. 2H and I).

**FIG 2.**
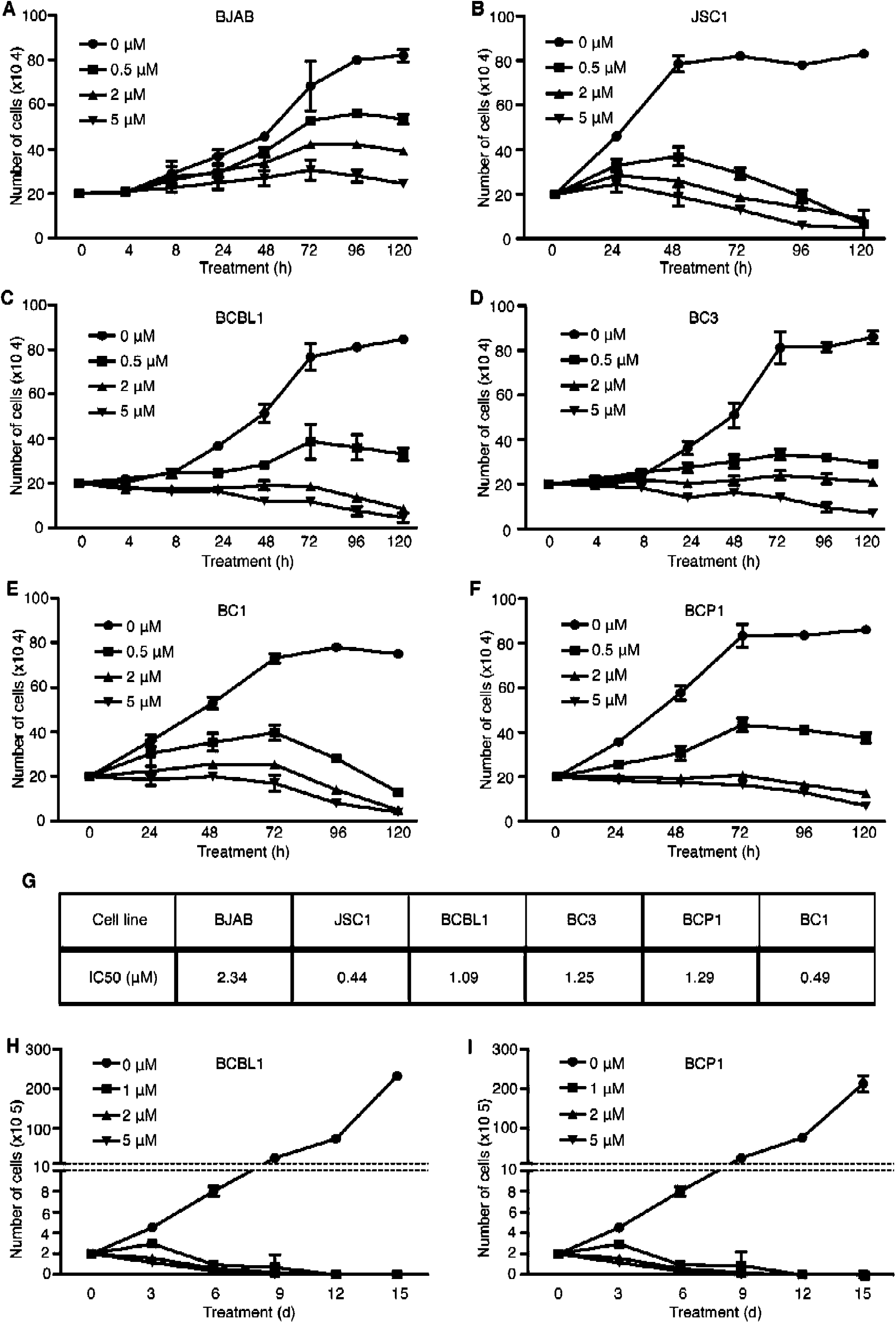
Cytarabine inhibits cell proliferation of primary effusion lymphoma cells. (A-F) Analysis of the cell proliferation of BJAB (A), JSC1 (B), BCBL1 (C), BC3 (D), BC1 (E) and BCP1 (F) cell lines treated with DMSO or different doses of cytarabine for 120 h. (G) IC50 (µM) of cytarabine for each PEL cell line. (H-I) Long-term effect of cytarabine on BCBL1 (H) and BC1 (I) cells over a period of 15 days. Cytarabine was replenished every 3 days.

### Cytarabine induces cell cycle arrest and apoptosis in PEL cells

To identify the mechanism of cytarabine-mediated cytotoxicity, we examined the effect of cytarabine on cell cycle. Cytarabine at 5 µM effectively induced cell cycle arrest in BCBL1 and BC3 cells atfter as short as 4 h of treatment (Fig. 3A). Similar result was observed with BJAB cells. Cytarabine at 5 µM induced apoptosis in 63% of BCBL1 cells and 49% of BC3 cells following 4 h treatment (Fig. 3B). Furthermore, treatment with 5 µM cytarabine for 24 h induced apoptosis markers including cleaved PARP1 (c-PARP1) and cleaved caspase 3 (c-caspase 3) in BCBL1, BC3, BC1, JSC1 and BCP1 cells (Fig. 3C). In contrast, cytarabine did not induce any apoptosis in BJAB cells (Fig. 3B and C).

**FIG 3.**
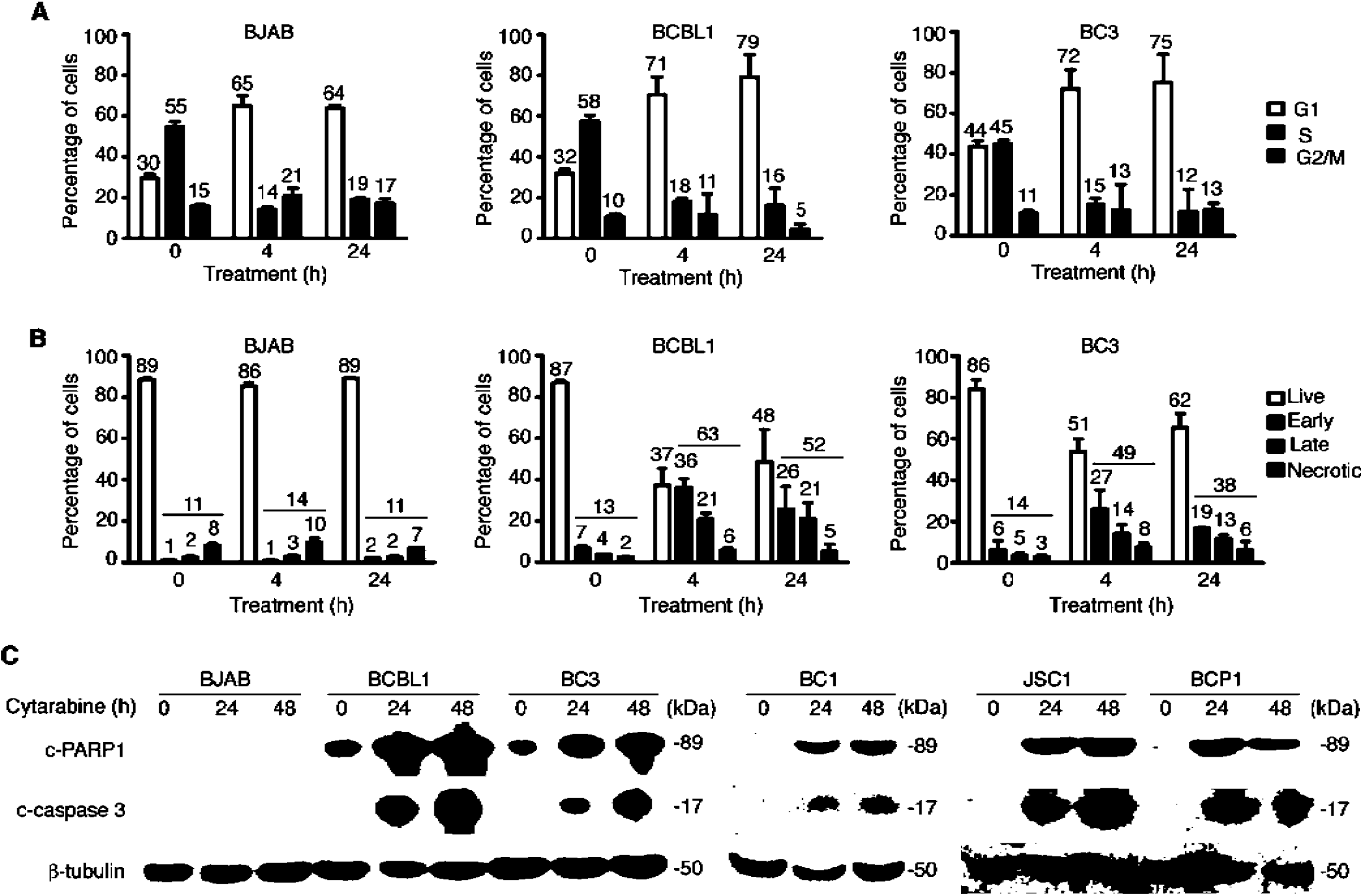
Cytarabine induces cell cycle arrest and apoptosis of primary effusion lymphoma cells. (A) Effect of cytarabine on cell cycle progression of BJAB, BCBL1 and BC3 cells. Cells treated with 5 μM cytarabine for 48 h were analyzed by flow cytometry after BrDu and PI staining. (B) Induction of apoptosis in BJAB, BCBL1 and BC3 cells by cytarabine. Cells treated with 5 μM cytarabine for 48 h were analyzed by flow cytometry after Annexin-5 and DAPI staining. Early apoptotic cells were those that were only positive for Annexin-5, late apoptotic cells were those that were only positive for DAPI, and necrotic cells were those that were positive for both Annexin-5 and DAPI. (C) Analysis of apoptosis markers cleaved PARP1 (c-PARP1) and cleaved caspase 3 (c-caspase 3) in BJAB, BCBL1, BC3, BC1, JSC1 and BCP1 cells treated with 5 μM cytarabine for 0, 24 and 48 h by Western-blotting.

### Cytarabine inhibits PEL initiation and progression, and regresses grown PEL

To examine the efficacy of cytarabine for PEL treatment, we employed a xenograft mouse model. We induced PEL in Nod/Scid mice by engrafting BCBL1-Luc cells. At day 3 post-engraftment, we treated the mice with either PBS or a liposome form of cytarabine Depocyt^®^ at 50 mg/kg every other day for 3 weeks. Mice treated with PBS started to gain weight at as early as 1 week as a result of PEL development while those treated with cytarabine maintained relatively constant weight (Fig. 4A). At week 3 post-treatment, we performed live bioluminescence imaging and detected strong signals in all mice treated with PBS (a-j in Fig. 4B and C). However, mice treated with cytarabine (k-t) had no detectable signal at this time point, indicating that cytarabine completely inhibited PEL growth.

**FIG 4.**
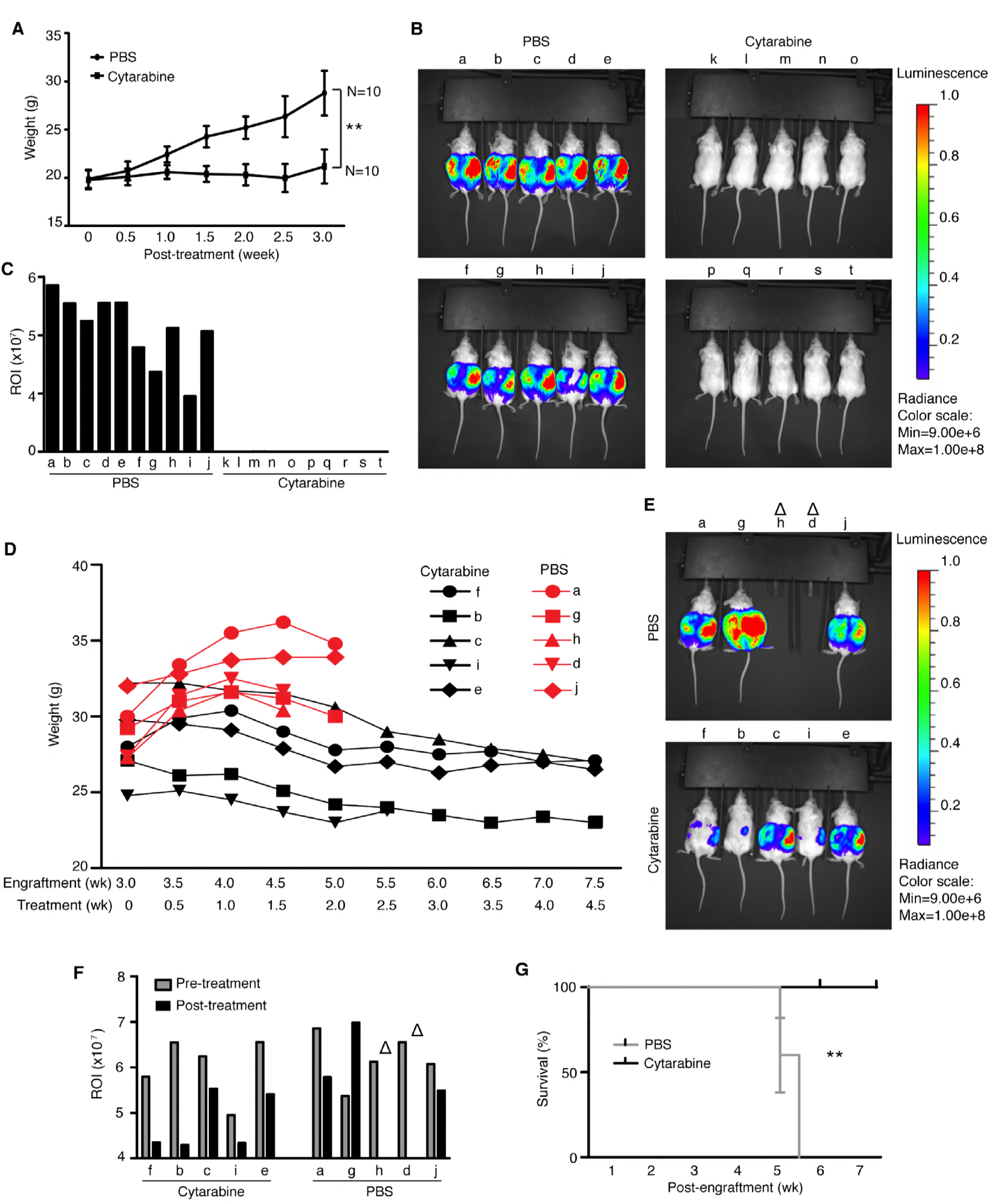
Cytarabine inhibits tumor initiation and progression in a xenograft mouse model of primary effusion lymphoma. (A) Weights of Nod/Scid mice intraperitoneally engrafted with 10^7^ BCBL1-Luc cells and treated 3 days later with a liposome form of Cytarabine (Depocyt®), every other days, for 3 weeks, were monitored twice a week. (B) Tumor burden of Nod/Scid mice described in (A) was analyzed by Luminescence assay at week 3 following the treatment of Depocyt^®^. (C) Luminescent signals from (B) were quantified and expressed in ROI (Total Radiant Efficiency [p/s]/[µW/cm^2^]). (D) Weights of Nod/Scid mice intraperitoneally engrafted with 10^7^ BCBL1-Luc for 3 weeks, and then treated with Depocyt®, every other days, for 4.5 weeks, were monitored twice a week. (E) Tumor burden of Nod/Scid mice described in (D) was analyzed by Luminescence assay at week 4.5 following the treatment of Depocyt®. “∆” symbol indicates mice euthanized before the Luminescence assay. (F) Luminescent signals from (E) were quantified and expressed in ROI. “∆” symbol indicates mice euthanized before the Luminescence assay. (G) Survival analysis of Nod/Scid mice described in (D).

Next, we examined if cytarabine could control or regress grown PEL. The mice engrafted with BCBL1-Luc cells for 3 weeks were randomly separated into two groups, and treated with PBS or Depocyt® at 50 mg/kg every other day for 4.5 weeks. Mice treated with cytarabine had reduced weights (f, b, c, i, e in Fig. 4D) indicating PEL regression while those treated with PBS continued to gain weight (a, g, h, d, j in Fig. 4D) indicating continuous PEL growth. At week 5 and 5.5 post-engraftment, mice “h, d” and “a, g, j” of the PBS group, respectively, died of PEL. Bioluminescence imaging at week 5 post-engraftment showed that the remaining 3 mice in the PBS group had strong luminescent signals (Fig. 4E and F). In contrast, the luminescent signals in all mice treated with cytarabine were dramatically reduced with those in mice “f, b, i” being reduced to almost undetectable levels, indicating that cytarabine effectively regressed most of the grown tumors (Fig. 4E and F). We compared the survival rates for both groups, and observed a statistically significant increase in survival rate for mice treated with cytarabine compared to mice treated with PBS (100% survival rate at week 8 post-engraftment for the cytarabine group *vs* 0% survival rate at week 6.5 for the PBS group) (Fig. 4G). These results demonstrated that cytarabine could be an effective drug for inhibiting PEL initiation and progression, and regressing grown PEL.

### Cytarabine inhibits KSHV latent replication

Since the survival of PEL cells depends on KSHV latent infection and multiple copies of viral episome, we determined if cytarabine might induce cytotoxicity by inhibiting KSHV latent infection. Treatment of BC1 and BCBL1 cells with cytarabine for 72 h decreased the level of intracellular KSHV DNA at least by half (Fig. 5A). Simultaneously, LANA transcript was reduced at least by half (Fig. 5B). Interestingly, LANA protein, which is essential for the replication and persistence of KSHV episome, was undetectable after 24 h of cytarabine treatment (Fig. 5C). Accordingly, <1 copy of KSHV genome per cell was detected after 3 days of cytarabine treatment compared to approximately 50 and 100 copies of KSHV genome per cell in the untreated BC1 and BCBL1 cells, respectively (Fig. 5D). Since cytarabine can be incorporated into RNA and DNA by competing with intracellular nucleotides (12, 13), we determined if it might directly inhibit KSHV latent replication and expression of KSHV latent genes. We pulsed BCBL1 cells with BrDu or 4sU for 4 h, and then immunoprecipitated the BrDu-labeled DNA or 4sU-labeled RNA to monitor *de novo* DNA or RNA synthesis, respectively. Whereas the amount of newly synthesized total DNA and RNA continued to increase over a period of 4 h in control cells treated with DMSO, those of cytarabine-treated cells did not increase, indicating that cytarabine inhibited the *de novo* syntheses of total DNA and RNA (Fig. 5E). By using qPCR for BrDu-labeled cellular DNA (18S and β-actin) and KSHV DNA (LANA), we detected inhibition of both cellular and KSHV DNA syntheses at as early as 30 min following cytarabine treatment (Fig. 5F). Interestingly, inhibition of β-actin and KSHV DNA syntheses seemed to be more efficient than 18S DNA synthesis. Similarly, by using RT-qPCR for 4sU-labeled cellular RNA and KSHV RNA, we detected inhibition of β-actin and LANA RNA syntheses by cytarabine (Fig. 5G). Inhibition of LANA RNA synthesis, which could be observed at as early as 15 min following cytarabine treatment, seemed to be more efficient than β-actin RNA synthesis. Interestingly, cytarabine treatment for up to 4 h had minimal effect on 18S RNA synthesis. Taken together, these results indicated that cytarabine inhibited the syntheses of cellular and KSHV DNA and RNA, which might account for its inhibitory effect on PEL cells and KSHV latent infection.

**FIG 5.**
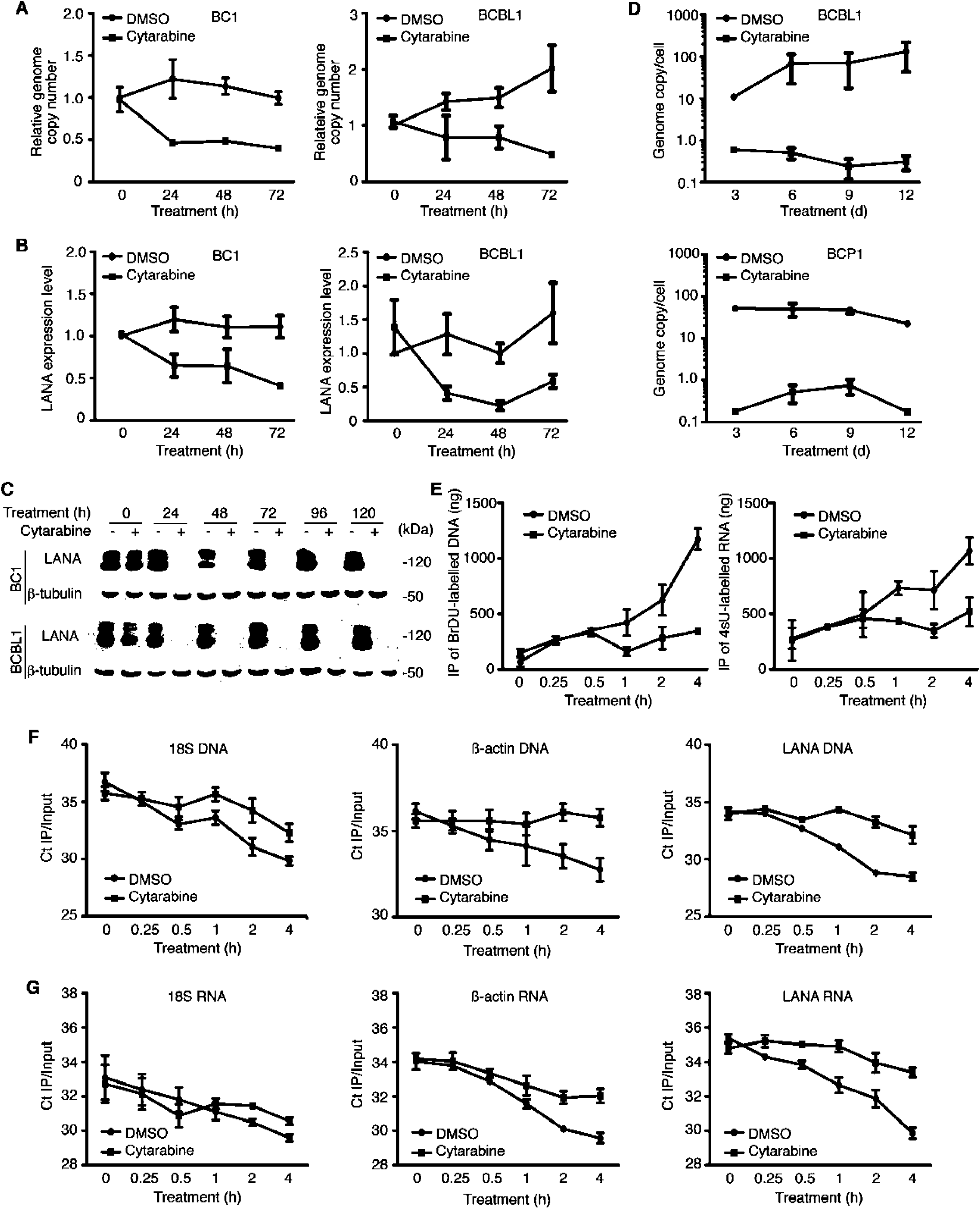
Cytarabine inhibits KSHV latent replication in primary effusion lymphoma. (A) Analysis of KSHV intracellular DNA in BC1 and BCBL1 cells treated with cytarabine by qPCR using LANA specific primers. (B) Analysis of the expression of LANA transcript in BC1 and BCBL1 cells treated with cytarabine by RT-qPCR using LANA specific primers. (C) Analysis of LANA protein in BC1 and BCBL1 cells treated with cytarabine by Western blot using a LANA specific antibody. (D) Quantification of KSHV genome copies per cell in BC1 and BCBL1 cells by qPCR after 12 days of cytarabine treatment. (E) Inhibition of DNA synthesis in BCBL1 cells treated with DMSO or cytarabine was analyzed by BrDU incorporation and quantification of immunoprecipitated BrDU-labeled DNA by Nanodrop. Inhibition of RNA synthesis in BCBL1 cells treated with DMSO or cytarabine was analyzed by 4sU incorporation and quantification of immunoprecipitated 4sU-labelled RNA by Nanodrop. (F) Inhibition of the syntheses of 18S, β-actin and LANA DNA in BCBL1 cells treated with DMSO or cytarabine was analyzed by qPCR on immunoprecipitated BrDU-labeled DNA from (E) using 18S, β-actin and LANA specific primers respectively. Results expressed in Ct were normalized with Ct of inputs. (G) Inhibition of the syntheses of 18S, β-actin and LANA RNAs in BCBL1 cells treated with DMSO or cytarabine was analyzed by RT-qPCR on immunoprecipitated 4sU-labelled RNA from (E) using 18S, β-actin and LANA specific primers respectively. Results expressed in Ct were normalized with Ct of inputs.

### Cytarabine inhibits KSHV lytic replication

Cell stress often triggers KSHV lytic replication and spread. Treatment of BCBL1 cells with cytarabine for 96 h did not increase KSHV lytic transcripts RTA, ORF59 and ORF65 (Fig. 6A), lytic proteins ORF-K8 and ORF65 (Fig. 6B), and virion production (Fig. 6C). While sodium butyrate (NaB) robustly induced KSHV lytic replication with increase of lytic transcripts RTA, ORF59 and ORF65 (Fig. 6A), lytic proteins ORF-K8 and ORF65 (Fig. 6B), and virion production (Fig. 6C), cytarabine completely inhibited this effect. These results indicated that cytarabine did not induce, rather it inhibited KSHV lytic replication.

**FIG 6.**
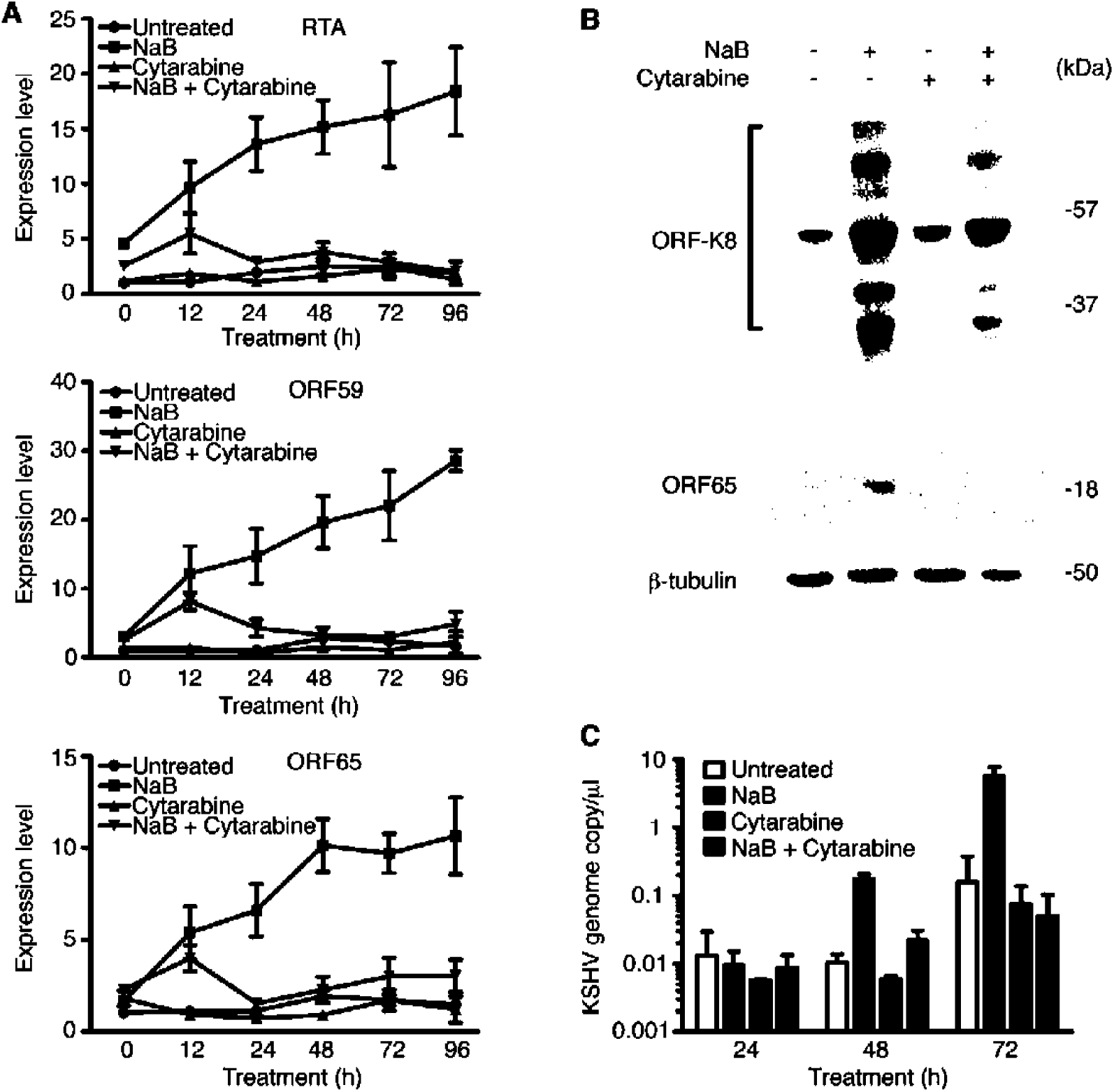
Cytarabine inhibits KSHV reactivation in primary effusion lymphoma. (A) Analysis of the expression of RTA, ORF59 and ORF65 transcripts in BCBL1 cells treated with DMSO, cytarabine, NaB or cytarabine + NaB by RT-qPCR. (B) Analysis of the expression of ORF-K8 and ORF65 proteins in BCBL1 cells treated with DMSO, cytarabine, NaB or cytarabine + NaB by Western blot using anti-ORF-K8 and anti-ORF65 antibodies. (C) Analysis of viral yield in BCBL1 cells treated with DMSO, cytarabine, NaB or cytarabine + NaB by PCR in supernatants pretreated with DNase.

## Discussion

There is currently no specific and efficient therapy for PEL. The common recommendation is “CHOP” chemotherapy, which is a combination of cyclophosphamide, doxorubicin, vincristine and prednisone (6). However, the outcome is dismal with a 1-year overall survival rate at 39.3% and an aggressive clinical course (9).

An anti-CD20 antibody has been developed for treating CD20+ B-cell NHLs. This approach can be considered for treating some rare cases of PEL expressing CD20 as it has shown some effect on MCD (14). Autologous stem cell transplantation in combination with high dose of chemotherapy is another strategy for treating PEL (15, 16). Finally, in HIV-infected PEL patients, targeting HIV infection and restoring immune functions by combined antiretroviral therapy (cART) alone has shown excellent outcomes (17). However, cART remains problematic because of potential adverse effects due to drug-drug interactions, which increase chemotherapy toxicities (18).

We have identified numerous “known” small molecules that inhibit the proliferation of KSHV-transformed cells but have minimal cytotoxicity to uninfected primary cells. These compounds are well characterized with most of them currently in clinical uses or trials, and hence are potential candidates for repurposing for KSHV-induced malignancies. Among them, cytarabine is effective in inducing cytotoxicity to PEL cells.

Cytarabine, or cytosine arabinoside, was first synthetized in 1959 and approved for clinical usages by FDA in 1969 (11). This drug has been used in therapy of several blood cancers such as AML, ALL and NHLs (11). Cytarabine is recently utilized for treating meningeal leukemias, lymphomas and recurrent embryonal brain tumors (19). Because of these multiple indications of treatment, the mechanism of action, which is mainly based on its inhibitory effect on DNA synthesis, the pharmacokinetics, and the toxicity of this compound in patients are well described (20). We have shown that cytarabine has a strong effect on PEL cells, inducing arrest and death *in vitro*, and completely abrogated tumor progression and regressed grown tumors in a PEL mouse model. These results indicate that cytarabine might be an ideal candidate for repurposing for PEL therapy.

Numerous studies have recently demonstrated that targeting pathways involved in differentiation and survival is a promising strategy for treating PEL. Activation of p53 with Nutlin-3 disrupted p53-MDM2 interaction and induced apoptosis of PEL cells and inhibits PEL progression (21). Silencing BLIM-1, a transcription factor involved in B-cell differentiation, led to PEL cell death (22). Triptolide inhibited cell proliferation and PEL progression by suppressing STAT3 activity, IL6 secretion and LANA expression (23). Chloroquine, an inhibitor of autophagy, induced a caspase-dependent apoptosis (24). Finally, thymidine analogue azidothymidine treatment (AZT), sensitized PEL cells to Fas-ligand and TRAIL-mediated apoptosis, and might be sufficient to restore T cell control *via* KSHV-specific T CD4+ response (25).

We have demonstrated that cytarabine induces cell cycle arrest and apoptosis in PEL cells by inhibiting DNA and RNA syntheses. Cytarabine can theoretically be incorporated into the DNA of any proliferating cells to induce DNA damage (12). It can also be incorporated into RNA to inhibit its polymerization, and therefore has an inhibitory effect on resting cells as well (13). In cells, cytarabine competes with the endogenous cytidine during nucleic acid synthesis after conversion into its triphosphate. Because the arabinose sugar of cytarabine sterically hinders the rotation of the molecule in DNA, it inhibits DNA replication. We have shown that cytarabine induces cell cycle arrest but not cell death in BJAB, an immortalized proliferating Burkitt’s lymphoma B cell line. This effect could be due to the incorporation of cytarabine into cellular DNA during S phase, resulting in the cytotoxicity on “normal” proliferating cells. However, in PEL cells, we have observed that cytarabine induces both cell cycle arrest and apoptosis, indicating alternative mechanism might be involved in its cytotoxic effect. Indeed, PEL cells are universally associated with KSHV infection and highly depend on KSHV latent proteins (3). We have shown that, in addition to cellular DNA and RNA, cytarabine inhibits KSHV DNA and RNA, resulting in the inhibition of replication of KSHV latent genome. This might explain the higher sensitivity of PEL cells to cytarabine than BJAB cells.

Nucleoside analogs are known to have efficient inhibitory effect on herpesviral persistent infections. By impairing the interaction between HSV-1 DNA and the cellular nucleosomes, cytarabine can inhibit HSV-1 replication (26). However, HBV is resistant to this drug but sensitive to its close relative adenosine analogue Ara-A (27). The chemically close fluoro-iodo-cytosine analogue FIAC is efficient against HSV-1, HSV-2 and EBV (28, 29). Because of its anti-tumor and anti-viral effects, it would be interesting to test if cytarabine is effective against cancers associated with other oncogenic viruses.

KSHV latently infected cells constitute a viral reservoir in the host. Upon stimulation by stress, inflammatory cytokines, calcium ionophores, phorbol ester or histone deacetylase inhibitors (Trichostatine A and NaB), KSHV can be reactivated from latency, expressing cascades of lytic genes and producing progeny virions (30–32). KSHV lytic replication in a small number of cells is essential for the spread and progression of early stage of KS (33). Numerous chemotherapies can induce reactivation of HSV-1 and EBV from latency, raising the concerns on these therapeutic approaches (34, 35). Our results show that cytarabine not only does not induce KSHV reactivation but also robustly inhibits the viral lytic program induced by NaB. In addition to KSHV, over 70% of PEL cases are co-infected by EBV, which itself is associated with several cancers (36). Similar to KSHV, EBV lytic replication participates in the spreading and pathogenesis of EBV-associated malignancies (37). Since cytarabine does not reactivate EBV from latency, it can be used for PEL treatment independent of the EBV status (38). Compared to other chemotherapies, the advantage of using cytarabine in PEL treatment is its multiple cellular and viral effects without increasing the spread of KSHV and tumor cells.

Taken together, we have identified cytarabine as an excellent candidate for reposition for treating PEL patients. It displays a strong inhibitory effect on PEL through multiple mechanisms. Since cytarabine is currently in clinical use and approved for treating other blood malignancies, it would be interesting to carry out advanced clinical trials to evaluate its usage and identify the optimal doses for treating PEL.

## Materials and Methods

### Cells culture

Rat primary embryonic mesenchymal stem cells (MM) and KSHV-transformed MM cells (KMM) were maintained in DMEM supplemented with 10% fetal bovine serum (FBS, Sigma-Aldrich), 4 mM L-glutamine, and 10 µg/ml penicillin and streptomycin (10). BJAB and PEL cell lines (JSC1, BCBL1, BC3, BC1 and BCP1) were maintained in RPMI-1640 supplemented with 20% FBS and antibiotics (39).

### High throughput screening

MM and KMM cells were seeded in 96 well plates at 5,000 cells/well for 16 h and then treated with small molecules at 5 µM final concentrations in 0.1% DMSO for 48 h. Cells treated with 0.1% DMSO and Bay11 were used as a negative control and a positive control, respectively. Cells were washed 2 times with 1X PBS and fixed with 4% paraformaldehyde for 15 min at room temperature prior to DAPI staining. Live cells were automatically counted with the Cellomics ArrayScan VTI HCS Reader (Thermo scientific) and the results were analyzed with the HCS Studio Cell Analysis Software (Thermo scientific). Number of live cells in DMSO control was set as 100% and used to normalize cells treated with different compounds. A total of 73 hits that gave at least 50% of cytotoxicity to KMM cells but less than 10% of cytotoxicity to MM cells were selected from the first round of screening, and then validated in a secondary screening, resulting in the selection of 50 final compounds. Secondary screening was carried out on ImageXpress Micro System (Molecular Devices) and the survival cells were automatically quantified using MetaXpress software (Molecular Devices). Compounds that showed effects similar to those of the first round screening with less than 20% variations were selected.

### Cell proliferation assay

MM, KMM and PEL cells plated at a density of 200,000 cells/well and treated by different reagents including DMSO, cytarabine or sodium butyrate (NaB) were counted daily using Malassez chamber.

### Cell cycle assay

Cell cycle was analyzed as previously described (39). PEL cells pulsed with 10 µM 5-bromo-2′-deoxyuridine (BrDu) (B5002, Sigma-Aldrich) were stained with propidium iodide (P4864, Sigma-Aldrich). BrdU was detected with a Pacific Blue-conjugated anti-BrdU antibody (B35129, Thermo Fisher Scientific).

### Apoptosis assay

Detection of apoptotic cells were carried out by staining the cells with DAPI and PE-Cyanine 7 conjugated anti-Annexin V antibody (25-8103-74, eBioscience) as previously described (39).

### Western blot

Western blotting was carried out as previously described (39). Primary antibodies to LANA (Abcam, Cambridge, MA); cleaved-PARP1 (CST); cleaved-caspase3 (CST); K8 (Santa Cruz); p53 (CST) and phospho-p53 (CST) were used. A monoclonal antibody to ORF65 was previously described (40).

### DNA extraction and qPCR

DNA extraction and quantitative real-time PCR (qPCR) were carried out as previously described using specific primers for β-actin (5’-TCCCTGGAGAAGAGGTAC-3’; 5’-AGCACTGTGTTGGCGTACAG-3’), 18S (5’-CAGCTTCCCAGAAACCAAAG-3’; ACCACCCATGGAATCAAGAA-3’) and LANA (5’-CCAGGAAGTCCCACAGTGTT-3’; AGACACAGGATGGGATGGAG-3’) (41). Relative gene expression levels were calculated using the 2-^∆∆^Ct formula and 18S as a loading control.

### RNA extraction and RT-qPCR

RNA extraction and reverse transcription qPCR (RT-qPCR) were carried out as previously described using specific primers for β-actin, 18S, LANA, RTA (5’-CACAAAAATGGCGCAAGATGA-3’; 5’-TGGTAGAGTTGGGCCTTCAGTT-3’), ORF59 (5’-CGAGTCTTCGCAAAAGGTTC-3’; 5’-AAGGGACCAACTGGTGTGAG-3) and ORF65 (5’-ATATGTCGCAGGCCGAATA-3’; CCACCCATCCTCCTCAGATA-3’) (41).

### De novo RNA synthesis

RNA was extracted from cells labeled with 500 µM of 4-thiouridine (4sU) (T4509, Sigma-Aldrich) for the indicated times in the presence of DMSO or 5 µM cytarabine in complete medium. RNA at 20 µg was biotinylated with 0.2 mg/ml of EZ-Link HPDP-Biotin (21341, Thermo Fisher Scientific) for 2 h at room temperature, and then subjected to phenol/chloroform extraction and isopropanol precipitation to remove the unlabeled HPDP-Biotin. The biotinylated-RNA fraction pellet was resuspended in 100 µl nuclease-free water and incubated with an equal volume of washed Dynabeads® MyOne™ Streptavidin C1 beads (65001, Thermo Fisher Scientific) for 30 min at room temperature. After 3 times of washing with washing buffer, the labeled RNA was eluded with 100 µl of 100 mM DTT, extracted by phenol/chloroform and precipitated with isopropanol. The final pellet was resuspended in 10 µl of nuclease-free water. cDNA was synthesized using 50 ng of RNA and specific primers of β-actin, 18S and LANA.

### De novo DNA synthesis

DNA was extracted from cells labeled with 10 µM BrdU (B5002, Sigma-Aldrich) for the indicated times in the presence of DMSO or 5 µM cytarabine in complete medium. DNA at 1 µg was resuspended in 100 µl nuclease-free water and fragmented with the DPNII restriction enzyme for 90 min at 37°C. Digested DNA was denaturated at 99°C for 10 min, cooled instantly in ice for 5 min and incubated with 5 µg of a biotinylated anti-BrdU antibody (ab2284, Abcam) in 100 µl of RIPA buffer at 4°C for 16 h. The Biotinylated-DNA was extracted with phenol/chloroform and precipitated with isopropanol. The biotinylated-DNA pellet was resuspended in 50 µl nuclease-free water and mixed with an equal volume of Dynabeads® MyOne™ Streptavidin C1 beads (65001, Thermo Fisher Scientific) in 50 µl of RIPA buffer for 2 h at 4°C. After 3 washes with washing buffer, the labeled DNA was eluded by boiling for 10 min in 100 µl 0.1% SDS solution. The DNA was extracted with phenol/chloroform, precipitated with isopropanol, and resuspended in 10 µl of nuclease-free water. The DNA was examined by qPCR with specific primers for β-actin, 18S and LANA.

### Animal experiment

For tumor initiation experiment, 20 female Nod/Scid mice at 5 weeks old were each intraperitoneally injected with 10^7^ BCBL1 cells expressing luciferase (BCBL1-Luc). At day 3 post-inoculation, mice were treated with PBS or a liposome form of cytarabine (Depocyt®) at 50 mg/kg every other day for 3 weeks and scaled for weight twice a week. At week 3 post-inoculation, mice were injected with Luciferin at 50 mg/kg and imaged with an IVIS Spectrum In Vivo Imaging System (Perkin Elmer). The signals were analyzed with the Living Image Software (Perkin Elmer) and expressed in ROI based on the [p/s]/[µW/cm²] formula.

For tumor regression experiment, 10 Nod/Scid mice each engrafted for 3 weeks with 10^7^ BCBL1-Luc cells were randomly split into 2 groups. One group treated with PBS and the other group treated with Depocyt® at 50 mg/kg every other day for 4.5 weeks were scaled for weight twice a week. At week 5 post-inoculation (*i.e*. week 2 post-treatment), live imaging was performed.

The protocols for the animal experiments were approved by the University of Southern California Institutional Animal Care and Use Committee under the protocol number #11722.

### Statistical analysis

Statistical analysis was performed using two-tailed t-test and P-value *P*≤0.05 was considered significant. Statistical symbols “*”, “**” and “***” represent P-values *P*≤0.05, ≤0.01 and ≤0.001, respectively, while “NS” indicates “not significant”. For survival study, Kaplan-Meier survival analysis was performed and statistical significance was calculated using the log-rank test.

## Acknowledgments

We thank members of Dr. Gao’s laboratory for technical assistances and helpful discussions. This work was in part supported by grants from NIH (CA096512, CA124332, CA132637, CA177377, CA213275, DE025465 and CA197153), and a grant from the University of Southern California Norris Comprehensive Cancer Center and The Choi Family Therapeutic Screening Facility to S-J Gao, and grants from NIH (CA082057, HL110609, DE023926, AI073099, AI116585) to J U Jung.

